# Normalized and Directional Interplay Scoring for the Interrogation of Proteoform Data

**DOI:** 10.1101/2024.11.18.624157

**Authors:** Karl F Poncha, Alyssa T. Paparella, Nicolas L. Young

## Abstract

Histone proteoforms, often presenting multiple co-occurring post-translational modifications (PTMs), are central to chromatin regulation and gene expression. A proteoform is a specific form of a protein that includes variations arising from genetic changes, alternative RNA splicing, proteolytic processing, and PTMs. Genomic context-dependent histone proteoforms define the histone code, influencing cellular phenotype by dictating interactions with DNA and chromatin-associated proteins. Understanding the dynamics of histone proteoforms is essential for elucidating chromatin-based regulatory mechanisms. Advances in middle-down and top-down proteomics methods enable accurate identification and quantitation of hundreds to thousands of proteoforms in a single run. However, the resulting data complexity presents significant challenges for analysis and visualization. Here, we introduce new computational methods to analyze the dynamics of histone PTMs and demonstrate their use in mouse organs during aging. We have developed and benchmarked two novel PTM crosstalk scores. The score that we term ‘Normalized Interplay’ addresses limitations of the original crosstalk score ‘Interplay’ providing a more complete and accurate measure of PTM crosstalk. The second score, ‘delta I’ or Directional Interplay is an asymmetric measure quantifying the magnitude and directionality of crosstalk between PTMs. Applying our two-stage scoring approach to data from CrosstalkDB, a community resource that curates proteoform-level data, reveals the dynamics of histone H3 modifications during aging. The source code is available under an Apache license at https://github.com/k-p4/ptm_interplay_scoring.

## Introduction

As mass spectrometry has become more powerful proteomics experiments increasingly encompass multiple levels of experimental and biological features. This includes analyzing proteomes from diverse tissues, using both technical and biological replicates, across various conditions and temporal stages. Such large-scale proteomics efforts yield insights into the system under study but pose significant challenges in data analysis and interpretation. These challenges are exacerbated when exploring protein post-translational modifications (PTMs). The proteoform represents the true physiological state of proteins, including post-translational modifications co-occur in combination on single molecules^1,2^. Multiple proteoforms of each protein in a sample exhibit unique sets of PTMs, each of which may present unique functional effects^3^. This complexity is particularly noticeable in hyper-modified proteins such as histones, where a vast array of histone proteoforms exist and are often interrelated in function^4–7^. The visualization of these highly interconnected systems of PTMs has not kept pace with data generation, hindering the extraction and presentation of functional and mechanistic biological insights. In this work, we introduce two new crosstalk scores to address the shortcomings of existing scores, including a new metric that quantifies the directionality of histone PTM crosstalk.

Histone proteins carry diverse epigenetic marks and play crucial roles in cellular processes like altering chromatin structure, genome maintenance, and gene regulation. Enzymatic PTM readers recognize specific modifications, recruiting PTM writers and erasers to modify other sites, a process influenced by existing PTMs. This interplay between PTMs is termed Crosstalk. Crosstalk can be either positive or negative and therefore we define this crosstalk as how the presence of one PTM can either potentiate or preclude the presence of another PTM as positive and negative crosstalk, respectively. One such canonical example is the binary mark of K9 methylation and S10 phosphorylation, which is an important mechanism in the progression of mitosis (a ‘methylation/phosphorylation switch’).

Accurately measuring PTM crosstalk is challenging, necessitating detailed information on PTM presence across multiple specific amino acids on a histone. Traditional bottom-up mass spectrometry typically analyzes 7-20 aa long oligopeptide chains, which most often does not capture this co-occurrence. Proteoform identification and quantitation is most directly achieved by top-down proteomics^4,6–10^. However, top-down approaches sometimes fail to distinguish isobaric PTM combinations from each other. Notably, the intact analysis of physiological histone H3 is challenging due to extensive isobaric species that are difficult to resolve chromatographically^11^. Middle-down mass spectrometry overcomes these limitations by analyzing longer histone segments such as the 1-50 aa N-terminal tail of Histone H3, enabling the assessment of proteoform abundances and the relationships between PTM combinations^12–15^.

Introduced in 2014, the interplay score was initially used to capture and quantify multilayered histone PTM interactions. While Interplay could only provide snapshots of identified PTM pairs under isolated conditions, Abundance Corrected Interplay (introduced in 2020) refined the interplay score to more precisely assess crosstalk, enabling comparisons across different conditions and temporal scales^16,17^. However, both suffer from similar limitations that we address here with the novel Normalized Interplay Score and Directional Interplay Score.

## Materials & Methods

### Data sources

Proteoform-level data used in this study was sourced from the CrosstalkDB database^17,18^. Specifically, we focused on datasets derived from histone H3 proteoforms quantitated from the brain, heart, liver, and kidneys of 3-, 5-, 10-, 18-, and 24-month-old C57BL/6 mice. The dataset is a rich source of PTM co-occurrences across different biological contexts. CrosstalkDB files CrDB000062 through CrDB000126 were downloaded in .CSV format for further processing. In addition to the aging dataset, we also applied our analysis to a dataset from the Kelleher lab, which used M4K – a method of total kinetic analysis to understand the mechanisms of how combinatorial methylation patterns are established on histone H3 K27 and K36 residues in myeloma cells expressing high (NTKO) or low (TKO) levels of MMSET/WHSC1/NSD2^19^. The dataset was chosen as the effective rate constants determined were used to infer the crosstalk and its direction between methylation states.

### Formulation of Normalized Interplay

#### Definitions

Given two PTMs, *PTM*1 and *PTM*2 their associated probabilities of occurrence are defined as follows:

- *P*_*PTM*1_ is the probability of occurrence of *PTM*1.
- *P*_*PTM*2_ is the probability of occurrence of *PTM*2.
- *P*_*PTM*1*PTM*2_ is the joint probability of co-occurrence of *PTM*1 and *PTM*2.

#### Prior Work: Interplay (I) Schwammle et al. (2014)

The Interplay score quantifies the relationship between PTM1 and PTM2, comparing their co-occurrence to what would be the expected co-occurrence if the two PTMs are independent, defined as:

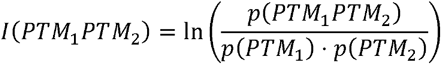

The log transformation ensures that:

- *I* > 0 when *PTM*_1_ and *PTM*_2_ co-occur more often than expected (positive association).
- *I* = 0 when *PTM*_1_ and *PTM*_2_ are independent.
- *I* < 0 when *PTM*_1_ and *PTM*_2_ co-occur less frequently than expected (negative association).

While the Interplay score provides insights into the association between PTMs, there are some limitations:

1. High negative association scores for low-frequency outcomes: The Interplay score tends to inflate association scores for PTM pairs involving low-frequency events. As the marginal probabilities *p*(*PTM*_1_) or *p*(*PTM*_2_) approach zero, the denominator becomes extremely small resulting in an Interplay score approaching negative infinity:

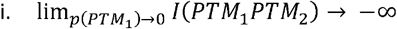
2. Low positive association scores for high-frequency outcomes: Conversely, Interplay tends to produce low association scores for pairs involving high-frequency outcomes due to the properties of the logarithmic function.
3. Unbounded values: The interplay score lacks fixed bounds in cases of perfect positive or negative association between two PTMs. This lack of consistent scale makes it difficult to compare interplay scores across multiple PTM pairs or datasets.

### Prior Work: Abundance Corrected Interplay (ACI) Kirsch et al. (2020)

Abundance Corrected Interplay (ACI) was introduced to address the limitations of the original Interplay Score. Each PTM’s abundance both relative to the other combination and the binary combination:

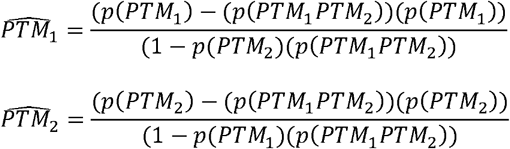

Yielding the ACI score as:

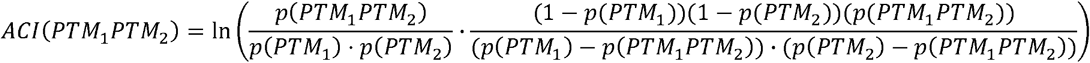

This formulation was designed to provide better symmetry between positive and negative interplay.

### Normalized Interplay (NI)

Since the limitations of the raw interplay score can affect interpretability, we introduce here the Normalized Interplay score. When two PTMs always co-occur, the probability of observing one given the presence of the other equals the joint probability of their co-occurrence. In this scenario the interplay score yields:

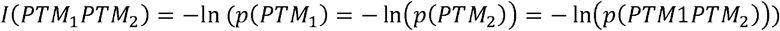

From which several options for normalization arise. The choice to normalize by the inverse natural logarithm of their joint probability is particularly appealing as it provides a symmetric normalization that adjusts both the upper and lower bounds:

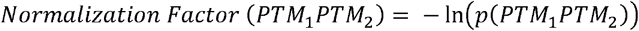

The normalization factor adjusts the Interplay score based on the ‘rarity’ of the co-occurrence of the two PTMs. Specifically, as *p*(*PTM*_1_*PTM*_2_) decreases, the value of ln(*p*(*PTM*_1_*PTM*_2_)) increases. The Interplay score is thus adjusted according to this scaling factor, such that more infrequent co-occurrences yield a higher (negative) adjustment:

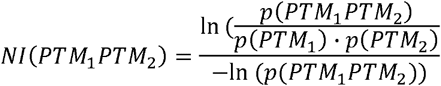

This normalization constrains the score within a bounded interval [− 1,1] offering a more intuitively interpretable metric:

### Formulation of the Directional Interplay Score (Δ*I*)

#### Rationale for Introducing Directionality in PTM Crosstalk Analysis

The interplay scores (I, ACI, and NI) discussed above treat the relationship between two PTMs as symmetric, assuming that the probability of PTM2 occurring given PTM1’s presence is equal to the probability of PTM1 occurring given PTM2’s presence. Mathematically, this is expressed as: *P*(*PTM*2|*PTM*1) - *P*(*PTM*1|*PTM*2), where *P*(*PTM*2|*PTM*1) represent the probability of PTM1 given PTM2 and vice-versa. However, this assumption overlooks that PTM crosstalk is often asymmetric, where one PTM exerts a stronger influence on the other in a non-reciprocal manner^20–23^. Symmetric measures of interplay collapse the distinct conditional probabilities *P*(*PTM*2|*PTM*1) und *P*(*PTM*1|*PTM*2), which introduces inaccuracies in interpretation, especially in hierarchical or sequential regulatory systems like PTM dependency networks, where directionality is critical for understanding regulatory interactions. To address this gap we introduce the directional interplay score which separates these conditional probabilities by independently evaluating *P*(*PTM*2|*PTM*1) *and P*(*PTM*1|*PTM*2).

#### PTM Information Categorization and Definitions

To derive the directional interplay score, we categorize the presence or absence of two PTMs, PTM1 and PTM2, into four terms:

- *a*: Joint probability where both PTM1 and PTM2 are present *(P(PTM1 ∩ PTM2))*
  ○ calculated as: abundance of the binary combination PTM1PTM2
- *b*: Joint probability where PTM1 is present and PTM2 is absent *(P(PTM1 ∩ P TM2))*
  ○ calculated as: normalized discrete abundance of *PTM1 − a*
- *c*: Joint probability where PTM2 is present and PTM1 is absent *(P(PTM2 ∩ P TM1))*
  ○ calculated as: normalized discrete abundance of *PTM2 − a*
- *d*: Joint probability where neither PTM1 nor PTM2 is present *(P(P TM1 ∩ P TM2))*
  ○ calculated as:
  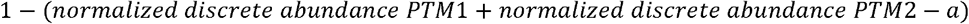

### Bayesian Framework for Directional Interplay

The directional interplay score (ΔI) leverages conditional probabilities to quantify the influence one PTM exerts on the other. By adopting a Bayesian framework, we treat the presence of PTM1 as information that updates our expectation of PTM2’s presence, akin to Bayesian updating^24^. For example, the conditional probability *P(PTM2*|*PTM1)*, representing the probability of PTM2 given PTM1, is computed as: 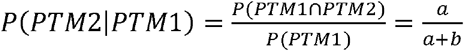

Here, *P(PTM2*|*PTM1)* is the posterior probability of PTM2 updating our prior knowledge of *P(PTM2)* based on the occurrence of PTM1. Similarly, the conditional probability of *P(PTM2*|*PTM1)* reflects the likelihood of PTM2 when PTM1 is absent:

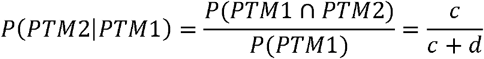

These conditional probabilities quantify how the presence or absence of PTM1 influences the likelihood of PTM2, and vice versa, allowing us to compute the directional interplay by comparing these probabilities.

### Formulation of the Directional Interplay Score (Δ*I*)

The directional interplay score ΔI measures the difference between the conditional probabilities of one PTM depending on the presence or absence of another. Thus, quantifying whether the presence or absence of one PTM increases or decreases the likelihood of observing the other PTM:

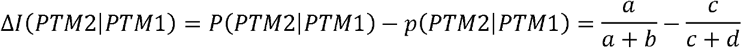

A positive value for ΔI(PTM2|PTM1) indicates that PTM2 is more likely to occur when PTM1 is present, suggesting a positive directional influence.

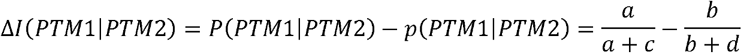

Similarly, this score quantifies how much PTM1 depends on the presence or absence of PTM2. A positive value for ΔI(PTM1∣PTM2) reflects a positive directional influence of PTM2 on PTM1. By independently evaluating *P(PTM2*|*PTM1) and P(PTM1*|*PTM2)*, ΔI captures the directional dependencies between PTMs, offering a more nuanced understanding of PTM crosstalk that cannot be quantified by symmetric measures.

### Data Processing Workflow

Jensen Aging dataset: Each run file from CrosstalkDB was divided into two H3 variant-specific files (H3.1/2 and H3.3). The percentage abundance of each proteoform was calculated by normalizing its intensity to the total intensity of the proteoform family, resulting in a per-family intensity of 100%. Proteoforms containing PTMs relevant to previous analyses were retained. The PTM code column was aggregated to full proteoform sequences, including unmodified residues. PTM combinations were generated using Python’s itertools.combinations function for all possible PTM configurations up to the proteoform level. Interplay scores were calculated for each binary PTM combination according to established formulae by mapping discrete PTM values to their constituent binary combinations. ΔI scores were computed by constructing contingency tables for each proteoform presence/absence pair. Normalized interplay scores were organized into matrices and visualized using the corrplot package in R. ΔI values were converted into source-target nodes for network analysis, with the antecedent as the source and the consequent as the target. Networks were constructed using Python’s NetworkX library for further analysis^25–27^.

Kelleher M4K dataset: The observed levels of all 15 detectable combinatorial methylations of the H3K27-K36 peptide in TKO and NTKO cells were downloaded from Table S1. ΔI scores were computed by constructing contingency tables for each PTM presence/absence pair in Excel (Microsoft Office 365 – Supplemental file 2).

## Results & Discussion

### Orientation Values for Interplay, Abundance Corrected Interplay, and Normalized Interplay

We examined the behavior of three PTM crosstalk scores across different scenarios of PTM co-occurrence (Summarized in Table 1).

**Table 1.** Contingency Table Illustrating the Co-occurrence of PTM1 and PTM2 Presence and Absence.

|  | *PTM2 (present)* | *¬PTM2 (absent)* |  |
| --- | --- | --- | --- |
| *PTM1 (present)* | *a* | *b* | *a+b* |
| *¬PTM1 (absent)* | *c* | *d* | *c+d* |
|  | *a+c* | *b+d* |  |

**Table 2.** Orientation values of PTM Crosstalk Scores.

| Condition of PTM co-occurrence | Interplay (I) | Abundance Corrected Interplay (ACI) | Normalized Interplay (NI) |
| --- | --- | --- | --- |
| Perfect positive crosstalk<br>(When two PTMs always co-occur) | $-\ln(P_{PTM_1PTM_2})$ | $+\infty$ | 1 |
| No crosstalk<br>(When two PTM co-occur as expected under independence) | 0 | 0 | 0 |
| Perfect negative crosstalk<br>(When two PTMs never co-occur) | $-\infty$ | $-\infty$ | -1 |

- Interplay: 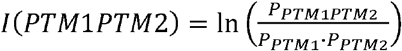
- Abundance Corrected Interplay: 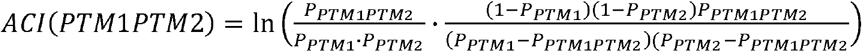
- Normalized Interplay: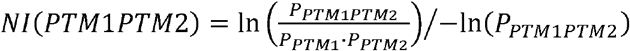

### Perfect Positive Crosstalk

In the scenario of perfect positive crosstalk, where PTM1 and PTM2 always co-occur. Here, as the joint probability *P*_*PTM*1*PTM*2_ = P_*PTM*1 = P*PTM*2_, the Interplay simplifies to − ln (*P*_*PTM*1*PTM*2_). For ACI, the correction factor introduces terms that adjust for the relative abundance of the PTMs. However, as the joint probability *P*_*PTM*1*PTM*2_ approaches *P*_*PTM*1_ or P_*PTM*2_, the denominator decreases towards zero, causing the ACI to diverge towards positive infinity: 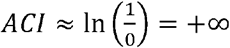. Since NI normalizes the interplay score by the negative logarithm of the joint probability: 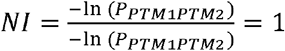.

### No Crosstalk

When PTMs occur independently, their joint probability equals the product of their individual probabilities: *p(PTM*_1_*PTM*_2_) = *p*(*PTM*_1_) · *p*(*PTM*_2_). In this case, all the scores essentially reduce down to ln*(1) = 0*, accurately capturing the scenario of no crosstalk.

### Perfect Negative Crosstalk

In the scenario of perfect negative crosstalk, where PTM1 and PTM2 never co-occur, the joint probability *p(PTM*_1_ *PTM*_2_) = 0. This results in: 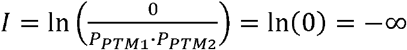. Similarly, for ACI, the numerator also becomes zero leading to ACI diverging to negative infinity. However, for NI as the joint probability tends towards zero, NI converges to its lower limit set to -1.

This bounded nature of NI ensures that even in cases of perfect positive or negative crosstalk, the score remains finite and interpretable, offering a stable measure of interplay.

### Functional Behavior of Interplay Scores Across Co-occurrence Probabilities

Each score responds uniquely to changes in the joint probability of PTM co-occurrence, influenced by the terms in their respective formulas. **Figures 1A and 1B** illustrate the behavior of the Interplay (I), Abundance Corrected Interplay (ACI), and Normalized Interplay (NI) scores across different co-occurrence probabilities for two PTMs. In Figure 1A, both I and ACI show non-linear responses. As the co-occurrence of PTMs approaches zero, both scores diverge towards negative infinity, indicating strong negative crosstalk. As their co-occurrence increases, they move towards zero and then ACI diverges towards positive infinity, signaling strong positive crosstalk.

**Figure 1.**
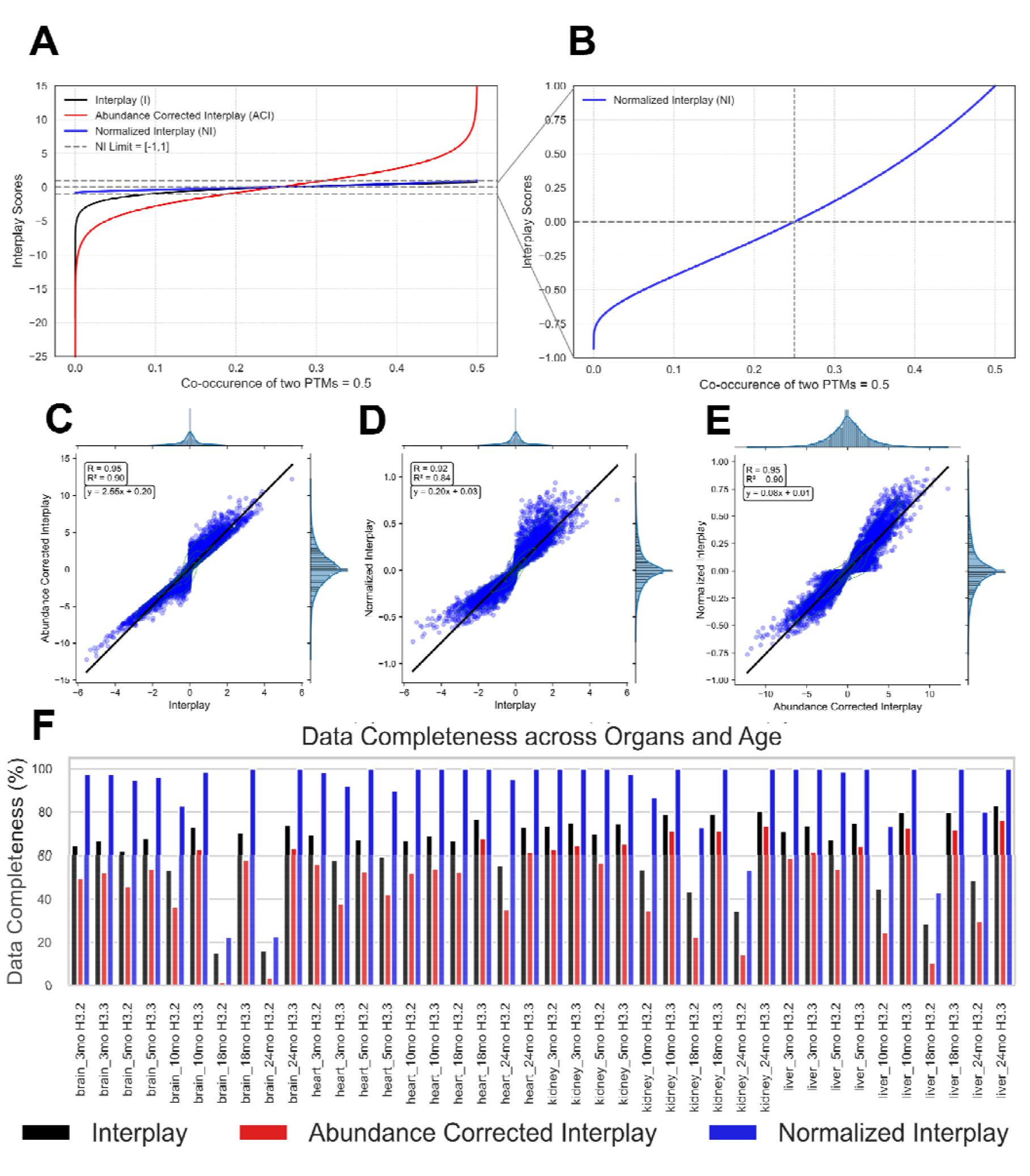
Comparison of the three Crosstalk scores: **(A)** The Interplay score (I), Abundance Corrected Interplay (ACI), and Normalized Interplay (NI) plotted as a function of the co-occurrence of two PTMs (y-axis) with respect to their frequency (x-axis). The ACI (red) adjusts for PTM abundance, while the NI (blue) normalizes interplay scores within the bounds of [- 1,1]. The original Interplay score (black) is shown for reference. **(B)** Detailed plot of Normalized Interplay (NI) highlighting the bounded range of [-1,1] as the co-occurrence of PTMs increases. This panel magnifies the NI transformation to clearly indicate how values differ at low co-occurrence rates. **(C-E)** Scatter plots of crosstalk between two PTMs quantified with each score across all organs, histones (H3.1/2 and H3.3) and ages. R^2^ denotes the coefficient of determination between each pair of plotted points. **(F)** Percentage of ‘Data Completeness’ considering all quantifiable instances of crosstalk across histones H3.1/2 and H3.3 from different organs and ages.

**Figure 1B** focuses on NI, which exhibits a linear and bounded response across the majority of the co-occurrence range. Unlike I and ACI, NI scales proportionally with changes in co-occurrence, avoiding overemphasis on minor fluctuations. This linearity ensures that NI remains stable and interpretable, providing a consistent measure that transitions smoothly from -1 to 1 as the co-occurrence of two PTMs moves from mutual exclusivity to perfect co-occurrence.

### Empirical Analysis of Interplay Scores in the Aging Dataset

Following the theoretical and functional comparison between the interplay scores We empirically analyze the Interplay (I), Abundance Corrected Interplay (ACI), and Normalized Interplay (NI) scores using the aging dataset, focusing on their pairwise correlations and data completeness across various organs and age groups.

### Correlation between scores

For PTM pairs where crosstalk is quantitated across all three scores, a high positive correlation is observed between all score comparisons. I and ACI exhibit a correlation of R^2^=0.90 (**Figure 1C**), indicating that while ACI follows the general trend of I, the deviations from the identity line show that ACI introduces specific abundance corrections that become more pronounced at extreme positive values. The comparison between I and NI shows a slightly lower correlation (R^2^=0.84 **Figure 1D**) reflecting NI’s normalization process. This, taken together with the high correlation between ACI and NI (R^2^=0.90, **Figure 1E**), indicates that NI incorporates the abundance corrections of ACI while introducing a normalization that differentiates its interpretation.

### Data Completeness Across Organs and Ages

Notably, a number of PTM pairs are not quantitated across all three scores and therefore not reflected in the correlation analysis between the scores. Across five organs and five ages, NI consistently demonstrates higher data completeness due to its ability to quantitate perfect positive and perfect negative crosstalk **(Figure 1F)**.

### Theoretical Bounds and Simulation of Delta I

#### Theoretical Bounds of ΔI

The directional dependency between two post-translational modifications (PTMs) is quantified by ΔI, which is bounded between -1 and 1. The extreme values of ΔI indicate specific types of relationships between PTM1 and PTM2. A ΔI(PTM2|PTM1) value of 1 corresponds to a perfect positive dependency, where PTM2 occurs exclusively when PTM1 is present, i.e., P(PTM2∣PTM1) = 1 and P(PTM2∣¬PTM1) = 0. Conversely, a ΔI(PTM2|PTM1) value of -1 indicates mutual exclusion, with P(PTM2∣PTM1) = 0 and P(PTM2∣¬PTM1) = 1, meaning PTM1 and PTM2 never co-occur. A ΔI(PTM2|PTM1) value of 0 reflects independence, where P(PTM2∣PTM1) = P(PTM2∣¬PTM1), signifying that the occurrence of PTM1 has no effect on the likelihood of PTM2.

#### Simulation of ΔI

To investigate the behavior of ΔI under different probabilistic conditions, we simulate values by fixing the probability of PTM1 (arbitrarily set at 0.5) and varying the conditional probabilities P(PTM2∣PTM1 and P(PTM2∣¬PTM1) across their full range from 0 to 1. This allows us to assess how various combinations influence the magnitude and sign of ΔI, capturing the directional dependency between PTM1 and PTM2. For each combination, the joint probabilities of PTM1 and PTM2 are computed and used to calculate ΔI, which quantifies the extent and direction of the dependency. The results are visualized in a 3D scatter plot (**Figure 2A**), where the x-axis represents P(PTM2∣PTM1), the y-axis represents P(PTM2∣¬PTM1), and the z-axis corresponds to the calculated ΔI values. The surface formed by the plot represents all possible theoretical values of ΔI, illustrating how the score smoothly transitions across varying conditional probabilities.

**Figure 2.**
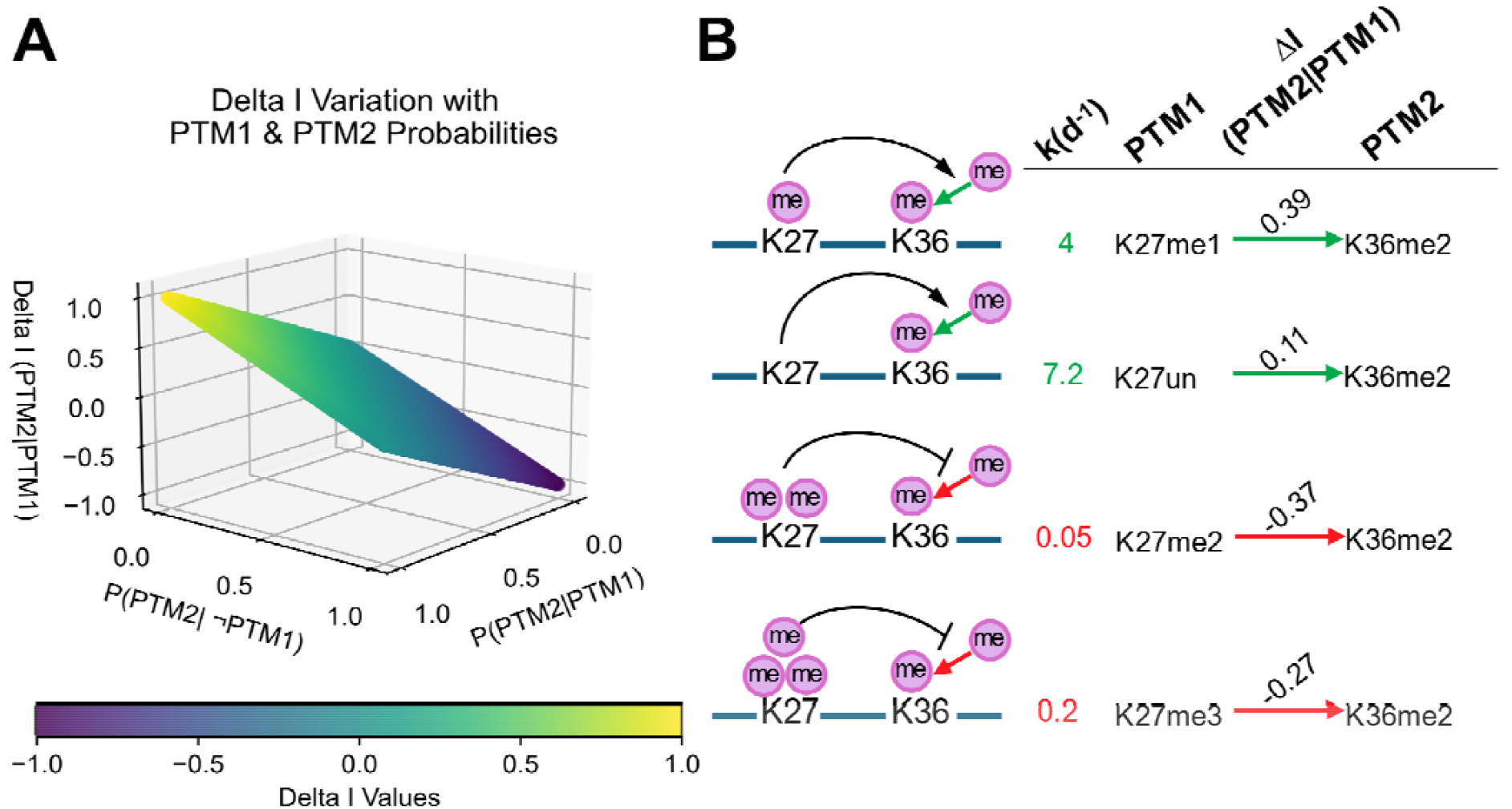
Simulation and benchmarking the directional Crosstalk score ΔI (delta I). Figure X: Simulation of ΔI variation and kinetic rate comparison for K27-K36 PTM crosstalk. **(A)** 3D Point cloud of theoretical ΔI (PTM2|PTM1) values, illustrating the relationship between conditional probabilities of two post-translational modifications (PTM1 and PTM2) and their influence on directional interplay. The x-axis represents the probability of PTM2 occurring given PTM1 P(PTM2∣PTM1), while the y-axis shows the reverse probability P(PTM1∣PTM2). The z-axis corresponds to the ΔI score, with a color gradient indicating the degree of crosstalk ranging from -1 (strong negative or antagonistic crosstalk -- purple) to +1 (strong positive or cooperative crosstalk -- yellow). This plot emphasizes how the likelihood of PTM1 influences the modification at PTM2 and vice versa, providing insight into the full range of directional interplay observed between modifications. This theoretical model offers a robust framework for quantifying PTM crosstalk under varying probability conditions. **(B)** Adapted from Figure S5C Zheng et al. (2012) Visual representation comparing directional interplay (ΔI) with experimentally derived kinetic rate constants for K27 and K36 methylation crosstalk. Each row represents a different methylation state of K27 (K27me1, K27un, K27me2, K27me3), highlighting how K27 modifications influence the dimethylation of K36 (K36me2). For each state, the kinetic rate constant k(d−1) is displayed alongside the corresponding directional ΔI score, with green arrows denoting positive crosstalk (synergistic PTM interaction) and red arrows indicating negative crosstalk (antagonistic interaction). ΔI values correlate with experimentally derived kinetic parameters to reflect the dynamic interplay between these modifications.

#### Crosstalk Between H3K27 and H3K36 Methylation States in TKO Cells

The relationship between H3K27 and H3K36 methylation has been extensively studied^19,28–31^. Kelleher et al.’s M4K kinetic model has provided detailed direct evidence of how methylation at one site influences the other. Their analysis of KMS11 TKO and NTKO cells revealed strong antagonism in the rate constants for the formation of dimethylated species when the other site is di- or trimethylated—a phenomenon they termed “bidirectional antagonism.” Conversely, unmethylated or monomethylated K36 facilitated faster methylation at K27 and vice versa.

Since no other directional scores exist for comparison, benchmarking the novel ΔI score is challenging. To demonstrate its utility, we applied ΔI scoring to the Kelleher dataset. The M4K model’s use of effective rate constants to infer directionality provides a distinct yet comparable framework for assessing how ΔI similarly captures the antagonistic interactions between H3K27 and H3K36 methylation states.

#### ΔI and M4K Analysis in TKO Cells

As shown in **Figure 2B**, when K27 is unmethylated, ΔI(K36me2|K27un) = 0.11, indicating that K36me2 is more likely to occur in the absence of K27 methylation. This positive crosstalk correlates with a high rate constant for the formation of K36me1→K36me2 (k = 7.2), reflecting a cooperative relationship where the lack of K27 methylation promotes K36 dimethylation. In the presence of K27 monomethylation, the directional crosstalk becomes even more pronounced, with ΔI(K36me2|K27me1) = 0.39, suggesting that K27me1 strongly favors the formation of K36me2. The rate constant for this transition, k = 4.0, supports this finding, indicating active promotion of K36 dimethylation by K27me1. As K27 progresses to higher methylation states, the interplay shifts towards antagonism. When K27 is dimethylated, ΔI(K36me2|K27me2) = -0.37, indicating that K27me2 hinders K36 dimethylation. This antagonism aligns with the much lower rate constant, k = 0.05, suggesting that K36 dimethylation is not favored in K27me2-containing cells. A similar trend is observed for K27 trimethylation, where ΔI(K36me2|K27me3) = -0.27 and the transition rate constant is k = 0.2, showing that K36 dimethylation is less likely when K27 is trimethylated.

The analysis shows that ΔI scores align closely with the M4K-derived effective rate constants, offering a quantitative reflection of the crosstalk between K27 and K36. ΔI provides a static measure of directional influence based on PTM abundance, indicating how the presence or absence of one methylation state impacts the likelihood of modification at the other. Positive ΔI values correlate with higher transition rates from M4K, reflecting cooperative crosstalk. Negative ΔI values correspond to lower transition rates, highlighting antagonism between the two sites.

M4K effective rate constants describe the rate at which specific PTM formations occur (e.g., K36me1 → K36me2), but these values do not necessarily reflect the final abundance of these methylation states. While a high rate constant indicates rapid formation, factors such as demethylation, further methylation to higher states, or other regulatory influences can impact the ultimate steady-state levels of these modifications. In contrast, the ΔI score captures the final directional dependency between PTMs based on their observed abundance, rather than the kinetic process leading to their formation. By quantifying this steady-state interplay, ΔI provides a static measure of the directional relationship between methylation states. Both metrics offer complementary perspectives: M4K provides insights into the dynamic formation rates of PTMs, while ΔI quantifies the stable outcome of their interactions. Overall, their agreement shows that ΔI is a valuable tool for analyzing PTM dependencies, especially when kinetic data is unavailable.

#### Normalized and Directional Interplay Scoring Histone Proteoforms During Aging

Normalized and directional interplay scoring delivers a more complete picture of the role of histone interplay in aging processes (**Figure 3**). During aging there is a progressive increase in positive crosstalk between K27me2 and K36me2 across multiple tissues and histone variants. In liver H3.2, the NI increases from 0.49 at 3 months to 0.57 at 5 months, reached 0.62 at 18 months, and climbed to 0.72 by 24 months. A similar pattern emerges in liver H3.3, where the NI increases from 0.45 at 3 months to 0.53 at 18 months and 0.57 by 24 months (**Supplemental File 1- S30-38**). Although the increase is more modest in brain H3.3, a comparable trend is evident during aging. Furthermore, these dynamics appear connected to acetylation states on histone tails, suggesting that these marks are associated with a more permissive and open chromatin structure. For instance, both K27me2 and H36me2 exhibit a moderately high NI of 0.11 with K14ac in the brain (**Supplemental File 1- S6-8)**.

**Figure 3.**
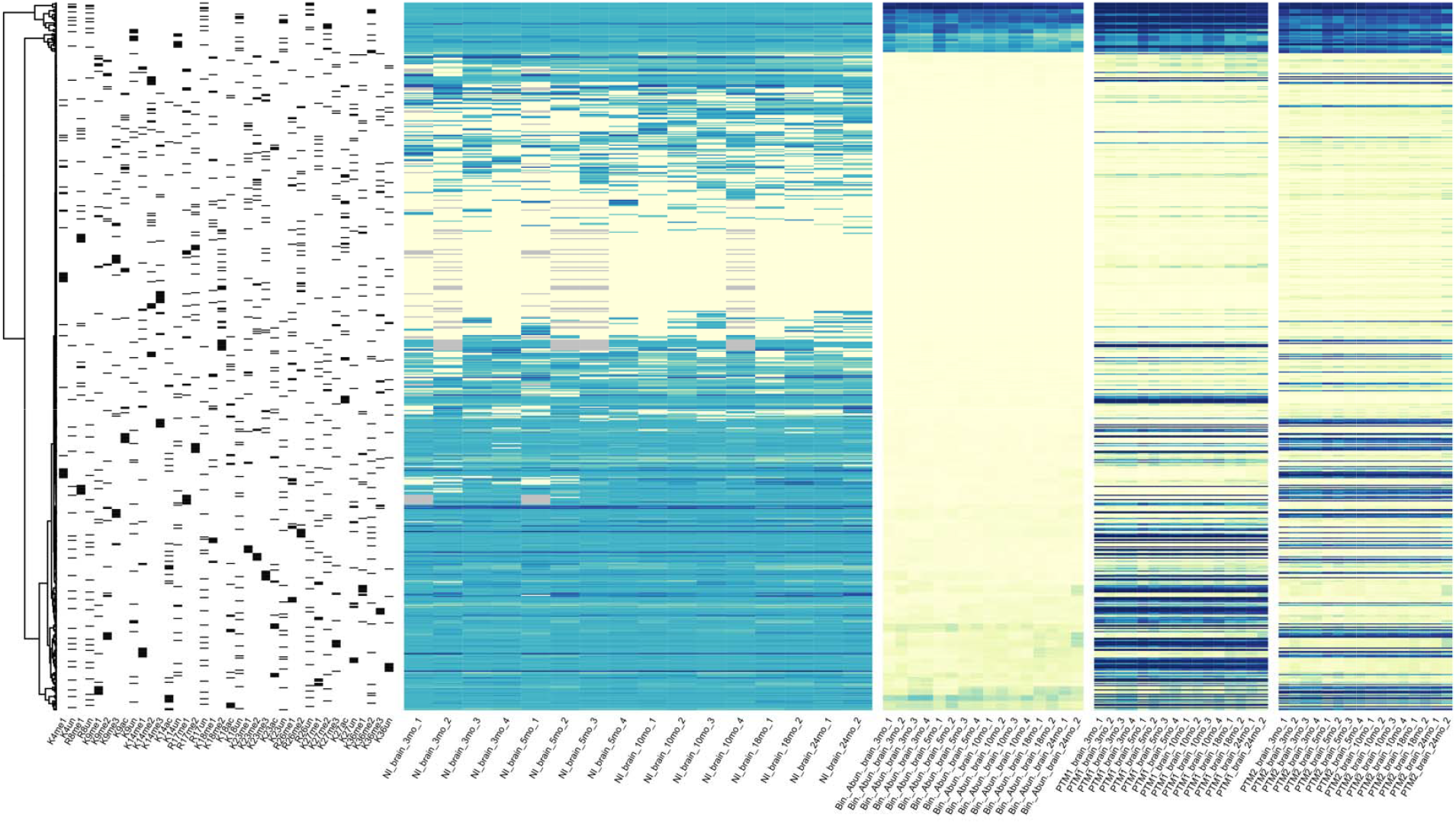
Heatmap Analysis of Post-Translational Modification Patterns and Interplay in H3.3 Histones from Brain Samples. This figure presents a complex heatmap of post-translational modification dynamics on H3.3 from brain samples, illustrating from left to right the presence of PTM combinations, normalized interplay (NI) scores, binary combinations, and the discrete abundances of PTM1 and PTM2 across different aging samples. Each heatmap is aligned to highlight relationships between different representations of PTM modifications. Samples are categorized by brain region, age, and biological replicate. **One-Hot PTM Heatmap (heatmap #1 -- leftmost):** This heatmap displays the presence (black) or absence (white) of specific PTM combinations across all brain samples using one-hot encoding. Each row represents a PTM combination, and each column corresponds to constituent PTM status. **Normalized Interplay Heatmap (heatmap #2):** This heatmap presents the normalized interplay (NI) scores for PTM combinations, quantifying the positive and negative crosstalk of PTMs within H3.3 histones. Higher NI scores (darker shades of blue) indicate stronger interactions or dependencies between modification states. **Binary Combination Heatmap (heatmap #3):** This heatmap shows the abundance of each PTM combination across brain samples. The color intensity (blue) corresponds to relative abundance, with darker shades indicating higher values. This representation reveals the percentage abundance of all PTM combinations. **PTM1 Abundance Heatmap (heatmap #4):** This heatmap displays the discrete abundance of the first PTM (PTM1) in each pairing across samples, with color intensity indicating relative abundance (darker shades represent higher levels). **PTM2 Abundance Heatmap (heatmap #5 --rightmost):** This heatmap illustrates the discrete abundance of the second PTM (PTM2) in each pairing across samples, with color intensity corresponding to relative abundance (darker shades indicate higher levels), similar to the PTM1 heatmap. All heatmaps are clustered based on the binary abundance of PTM combinations, revealing potential patterns and relationships between samples. Column names reflect specific sample characteristics, including brain region, age, and biological replicate. The heatmaps are scaled to emphasize differences in interplay and abundance across samples, offering insights into the combinatorial nature of PTMs in H3.3 histones during brain development and aging.

In liver H3.2, K9acK36me2 shows a positive NI of 0.38, and in liver H3.3, an NI of 0.13. These findings support the idea that K36me2 has a strong relationship with acetylation, contributing to the formation of permissive chromatin regions that are strengthened with age (**Supplemental File 1- S35**). Similarly, the crosstalk between K9me1 and K27me2 shows an upward trend, though with some variation across tissues and histone variants. In liver H3.2, the score starts at 0.46 at 3 months, peaks at 0.55 at 5 months, and then slightly declines to 0.47 by 18 months, remaining relatively stable at 0.45 at 24 months. This pattern suggests an early peak in positive crosstalk between these modifications, followed by a stabilization as the organism ages. In brain H3.2, the score increases steadily from 0.25 at 3 months to 0.34 by 18 months and continues to increase to 0.44 by 24 months, indicating a more sustained growth in positive interplay with age in neural tissues, where dynamic transcriptional regulation is critical (**Supplemental File 1- S5, 6**).

Several PTM pairs demonstrate increasingly negative crosstalk with age, reflecting growing mutual exclusion between modifications that mark distinct chromatin states. One of the most striking examples is the interplay between K9me2 and K27me2, which becomes progressively more negative as the organism ages. In liver H3.2, the score starts at -0.70 at 3 months, decreases to -0.88 by 5 months, and further drops to -0.91 at 18 months and -0.95 at 24 months. A similar trend is observed in liver H3.3, where the interplay score becomes more negative from -0.51 at 3 months to -0.66 by 24 months. This progressive decrease suggests that K9me2 and K27me2 become increasingly segregated, likely marking distinct repressive and permissive chromatin domains with greater precision as liver tissue ages. There is a less marked albeit consistently negative crosstalk between K9me2 and K27me2 in the brain consistently remaining at ∼-0.48 across ages. Indicating that K9me2 and K27me2K36me2 mark non-overlapping chromatin regions that are functionally distinct, repressive vs more permissive chromatin.

Interplay analysis also reveals organ-specific dynamics of aging. For instance, the mark K9me1K27me1 exhibits an interesting relationship. From 3 to 18 months, the crosstalk between K9me1 and K27me1 in heart tissue shows a clear trend of decreasing mutual exclusion. At 3 months, the average NI is strongly negative at - 0.82, indicating that these modifications are mostly mutually exclusive, marking distinct repressive chromatin regions. This exclusivity continues at 5 months, with a consistent NI of -1, showing that K9me1 and K27me1 rarely co-occur in the same regions. However, by 10 months, the mutual exclusion starts to weaken, as reflected by the average NI of -0.55. This change indicates that some chromatin regions start to show minor co-occurrence of K9me1 and K27me1, though exclusion still dominates. At 18 months, this trend becomes more pronounced, with the average NI shifting to 0.14, signaling a significant increase in co-occurrence. By this stage, K9me1 and K27me1 are no longer strictly exclusive, and they begin to mark overlapping chromatin regions, suggesting an evolving chromatin structure in aging heart tissue (**Supplemental File 1- S11-19**).

### The value and limitations of interplay scoring

The unique value of top- and middle-down proteomics is the ability to quantitatively study proteoform biology; however, effective use of proteoform information to derive fundamental mechanisms requires novel data analytics approaches. Recent work has shown that proteoform biology does not primarily operate at individual PTMs or single proteoforms but a regime between these extremes where proteoform families or themes of combinatorial PTM interplay dominate. Like PTMs, individual proteoforms mostly do not have an injective functional relationship. Thus, it is not sufficient to quantitate proteoforms and connect them to function. Rather, we must endeavor to understand the interplay between combinatorial PTMs. This requires unique data analytics and visualization of the high dimensional combinatorial data of proteoforms to identify functional units shrouded within this data. Prior work has provided useful tools to derive this important biology; however, prior scores have several limitations and lack clear intuitive interpretation. We propose here an improved method, Normalized Interplay, to improve symmetry between synergistic and antagonistic interplay and provide scores that are bounded. The resulting scoring function produces output that enables comparison between the strength of negative and positive interplay and intuitive bounds, where 1 represents perfect synergy and -1 represents perfect antagonism. These bounds also enhance data completeness because perfect synergy and antagonism are relatively common. Interplay is not limited to two PTMs and thus the measured relationship between two PTMs is likely influenced by other PTMs and the overall physiological state of the cell. The framework we present here is readily extended to higher dimension combinations, although we focus here on binary combinations. Binary combinatorial interplay is not limited to mutual co-occurrence and directionality of these relationships is essential to understanding proteoform mechanisms and can be readily inferred from quantitative proteoform data. We address this here with the introduction of Directional Interplay Score (Δ*1*). This scoring function provides the first systematic method to quantify the directionality of these relationships directly from proteoform data. The order of operations is fundamental to revealing proteoform mechanisms. There are no known mechanisms where the writing or erasing of multiple PTMs is truly concomitant. Thus, there is no reason to expect that the biochemistry of a reversed order of operations should be similar. Indeed, hierarchical highly directional relationships are abundant in the currently limited proteoform biology literature.

## Conclusion

We present here two complementary novel scoring methods for the elucidation of proteoform biology directly from quantitative proteoform data. The source code for these algorithms is made available to the top-down proteomics community. These methods are convenient and intuitive for the deeper inquiry into proteoform data. It is important to recognize that, like any discovery tool, these scores only infer mechanistic relationships. Full elucidation of mechanism requires the positing and testing of a hypothesis often with complementary approaches^32,33^. The methods presented are most useful for discovery of hidden functional relationships within quantitative proteoform data. However, the methods presented here can also be used effectively to test hypotheses when experiments are carefully designed for rigor.

## Acknowledgments

This work was supported by Howard Hughes Medical Institute grant to A.T.P: GT16569. This work was supported by the National Institutes of Health grants to NLY: R01GM139295, P01AG066606, R01CA193235, R01AG074540, R01CA276663, R01AG085751, and R01NS136375. The authors declare no conflicts of interest.

