## Supplemental File 1 for "Normalized and Directional Interplay Scoring for the Interrogation of Proteoform Data"

Normalized Interplay Matrices

Each matrix consists of each PTM pair represented on the axes and their intersection colored by their Normalized Interplay Score (ranging from -1 to +1). Quantitated across the Jensen Aging Dataset.

### Figure S1

Brain, 10 months, Histone H3.2

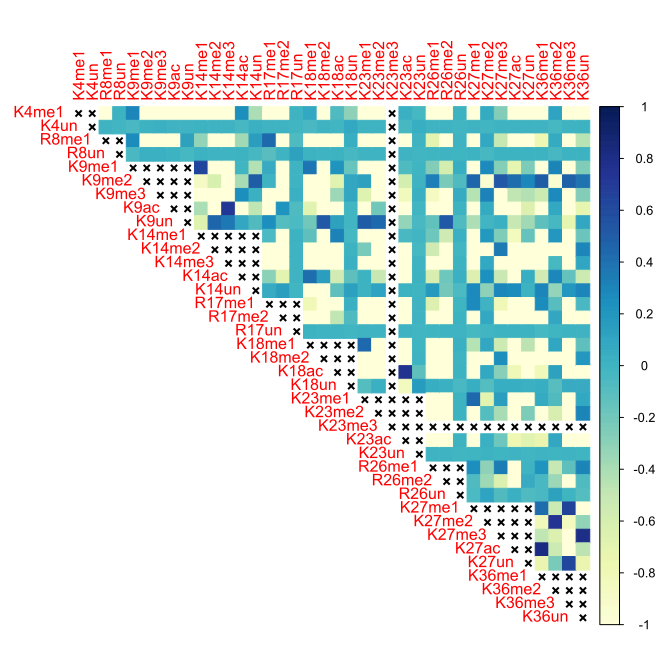

### Figure S2

Brain, 10 months, Histone H3.3

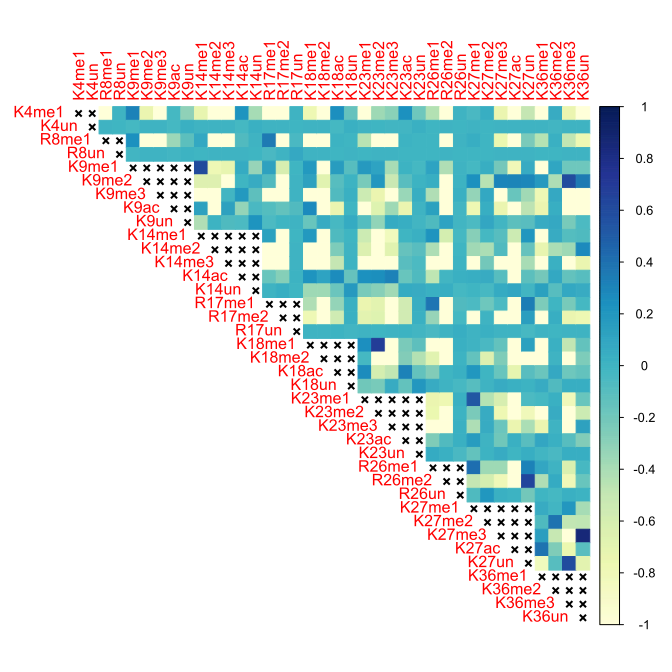

### Figure S3

Brain, 18 months, Histone H3.2

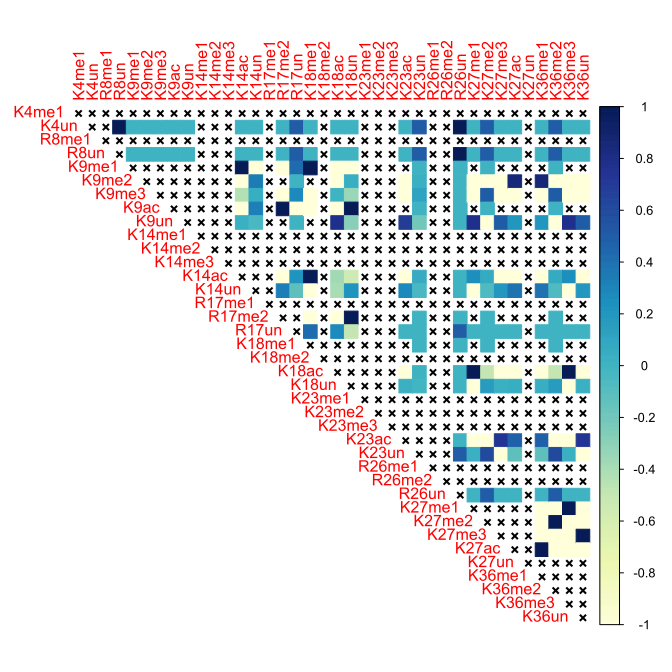

### Figure S4

Brain, 18 months, Histone H3.3

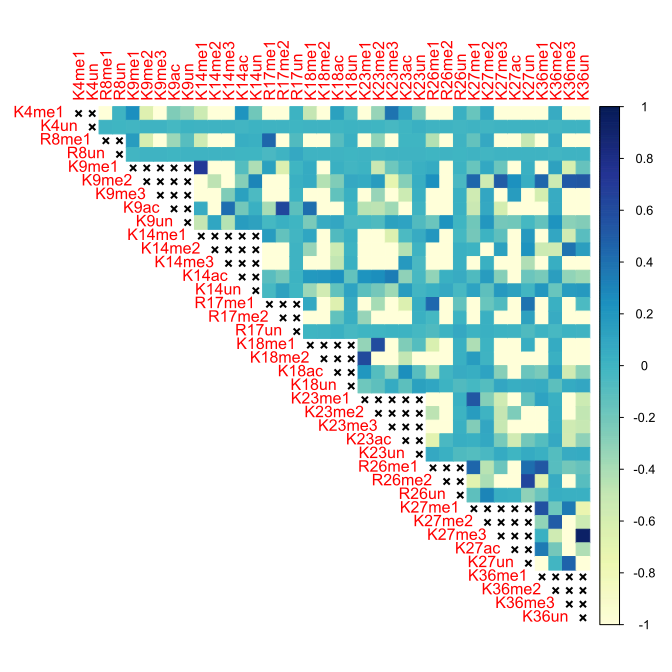

### Figure S5

Brain, 24 months, Histone H3.2

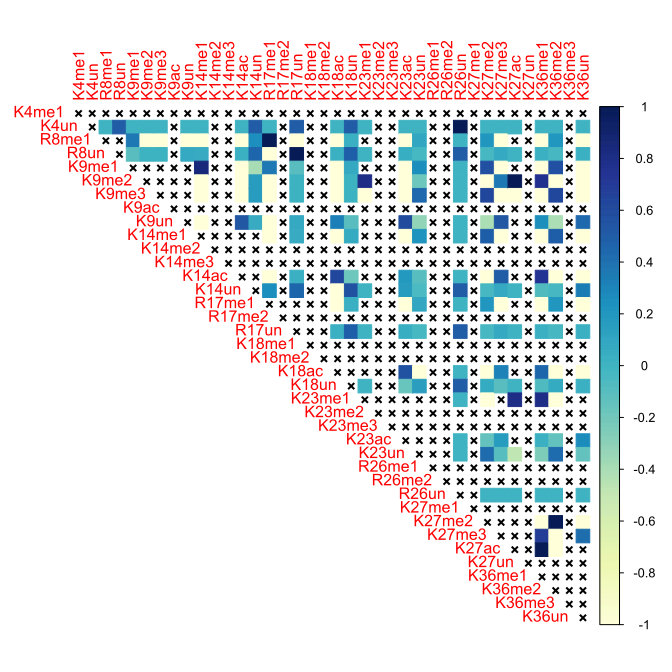

### Figure S6

Brain, 24 months, Histone H3.3

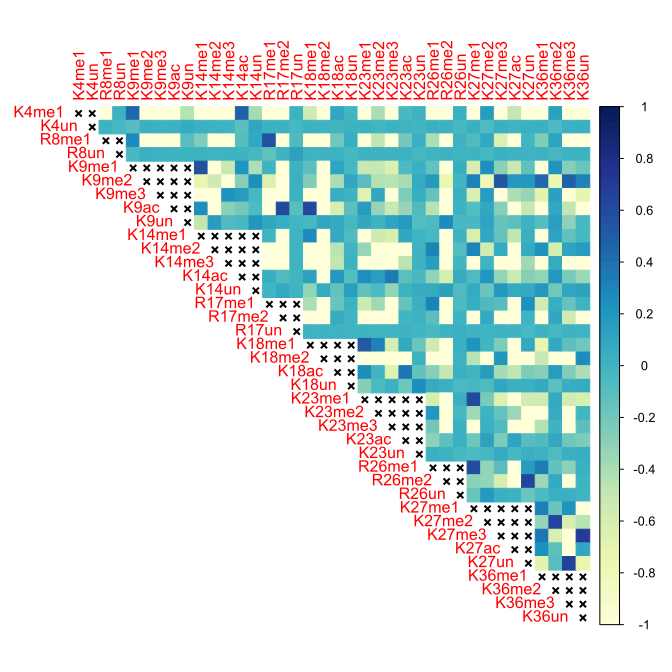

### Figure S7

Brain, 3 months, Histone H3.2

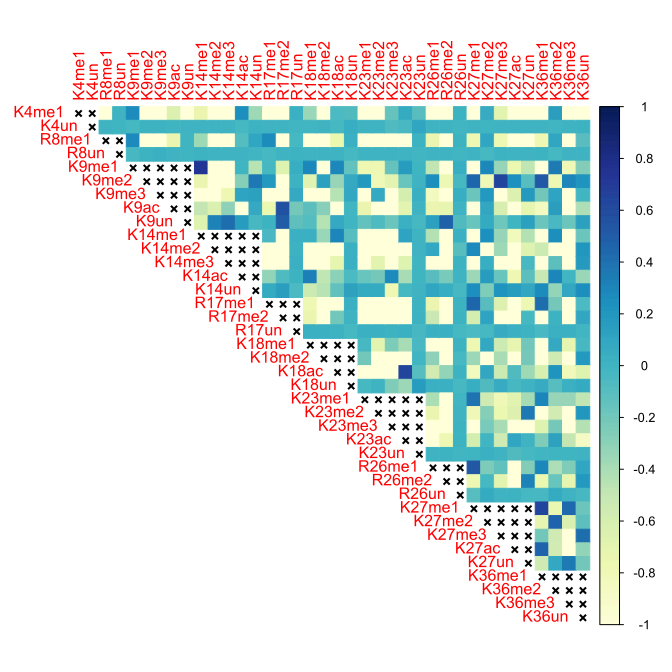

### Figure S8

Brain, 3 months, Histone H3.3

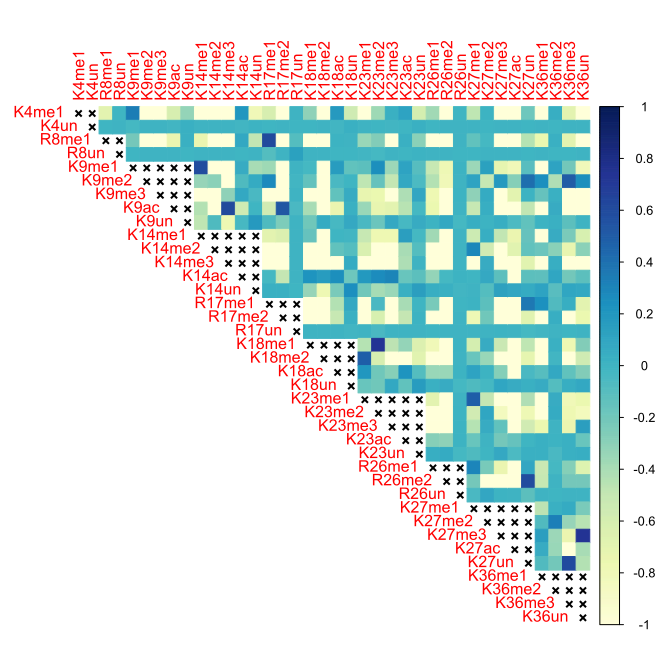

### Figure S9

Brain, 5 months, Histone H3.2

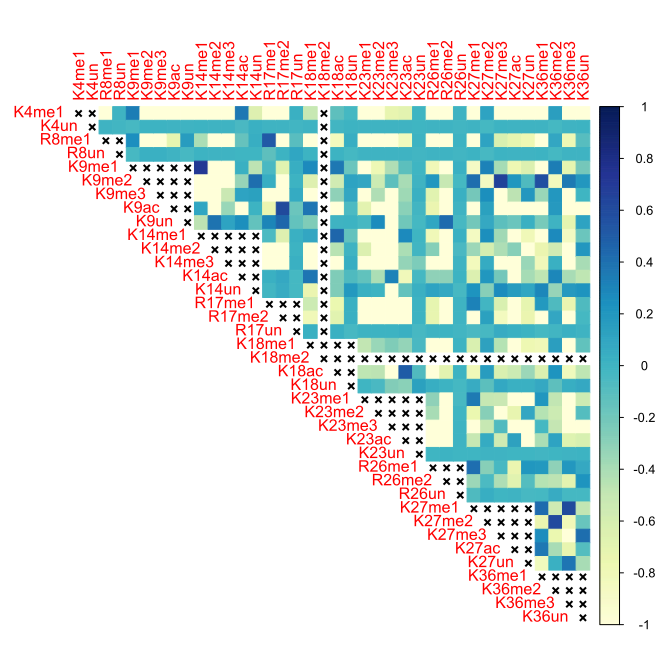

### Figure S10

Brain, 5 months, Histone H3.3

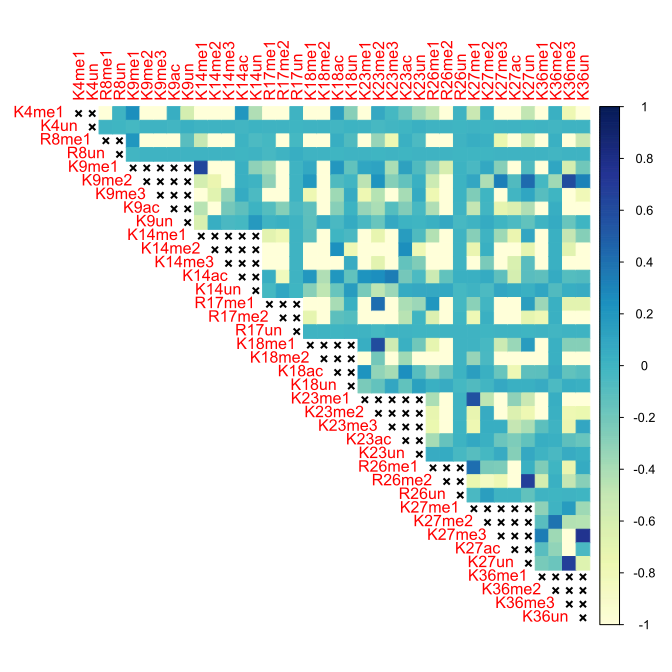

### Figure S11

Heart, 10 months, Histone H3.2

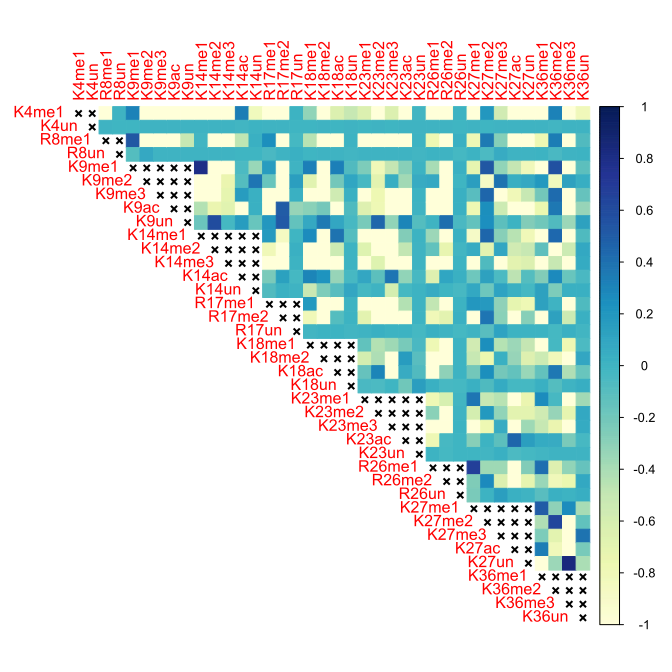

### Figure S12

Heart, 10 months, Histone H3.3

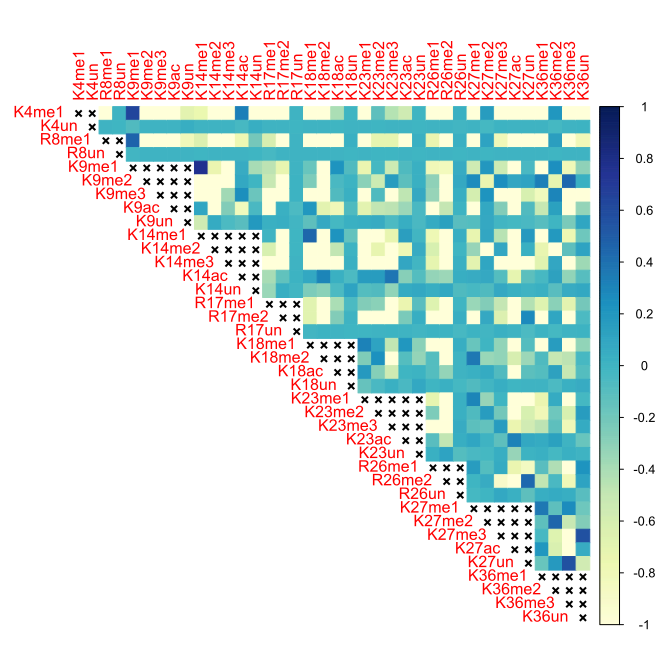

### Figure S13

Heart, 18 months, Histone H3.3

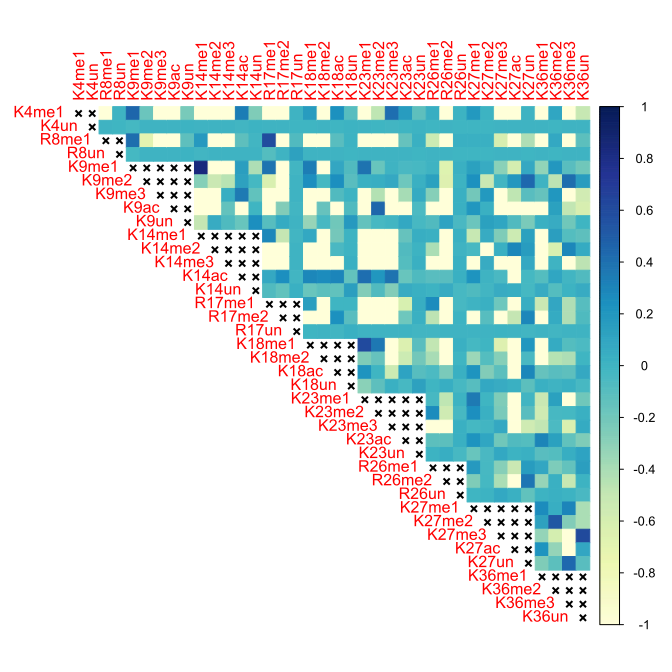

### Figure S14

Heart, 24 months, Histone H3.2

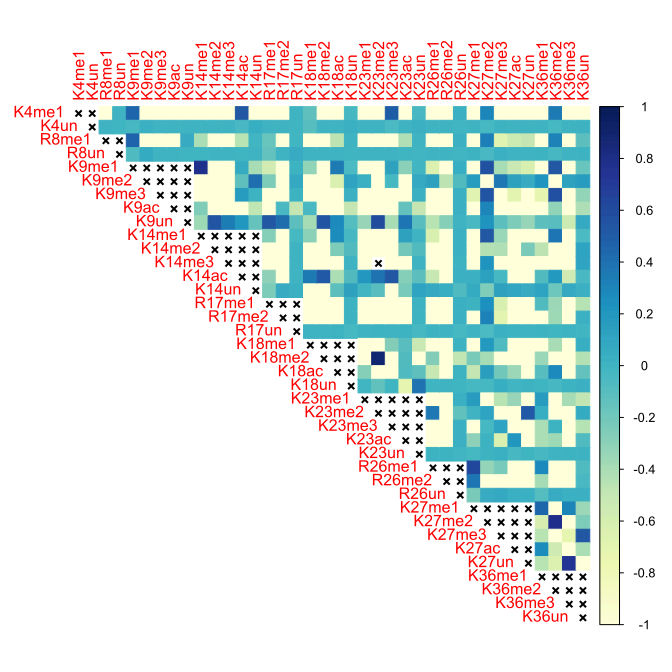

### Figure S15

Heart, 24 months, Histone H3.3

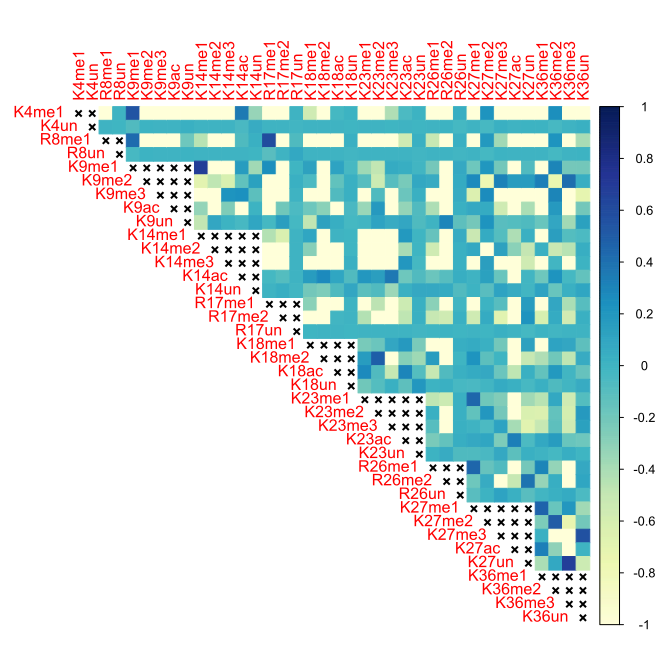

### Figure S16

Heart, 3 months, Histone H3.2

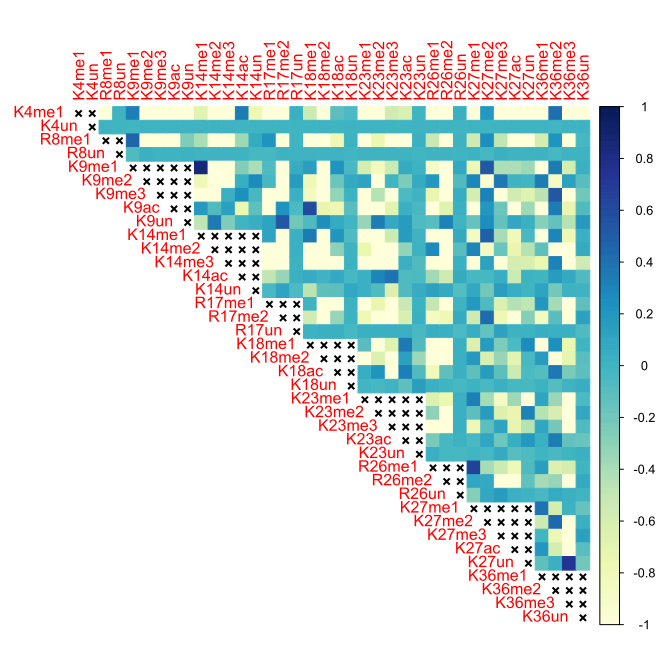

### Figure S17

Heart, 3 months, Histone H3.3

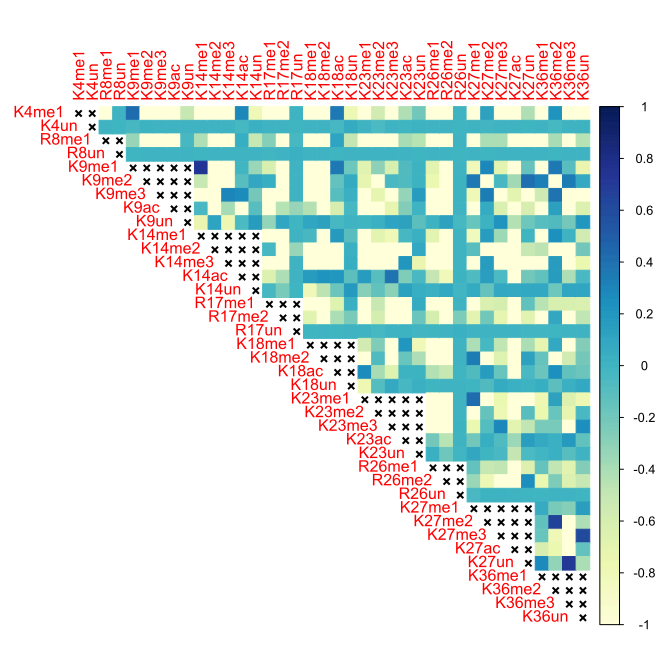

### Figure S18

Heart, 5 months, Histone H3.2

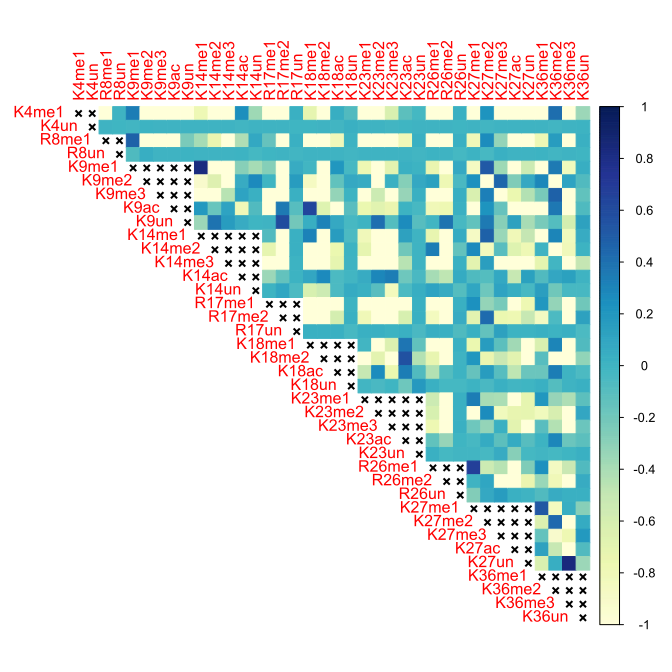

### Figure S19

Heart, 5 months, Histone H3.3

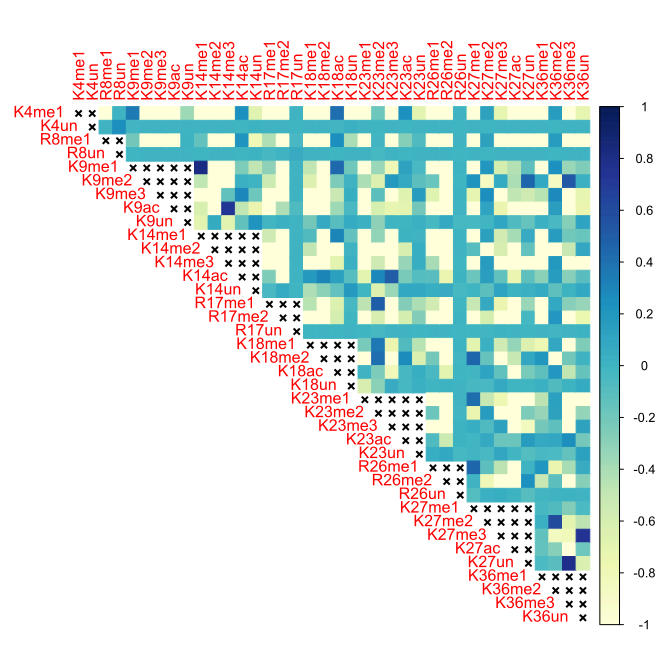

### Figure S20

Kidney, 10 months, Histone H3.2

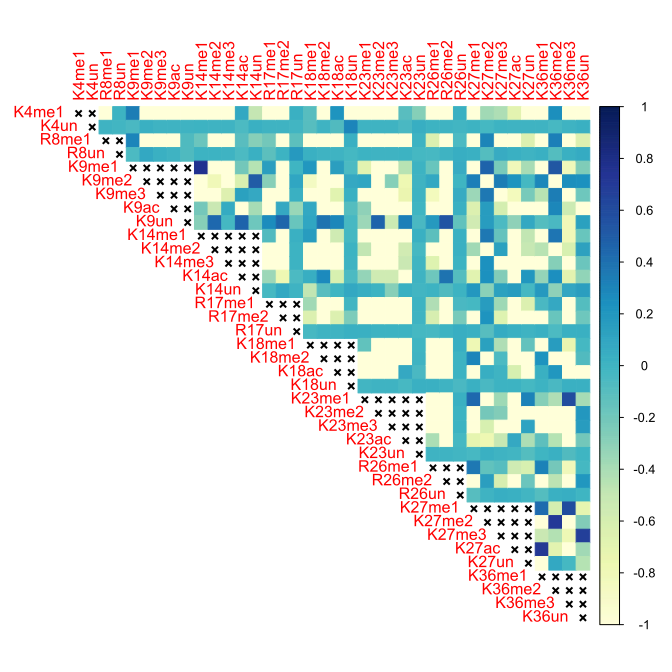

### Figure S21

Kidney, 10 months, Histone H3.3

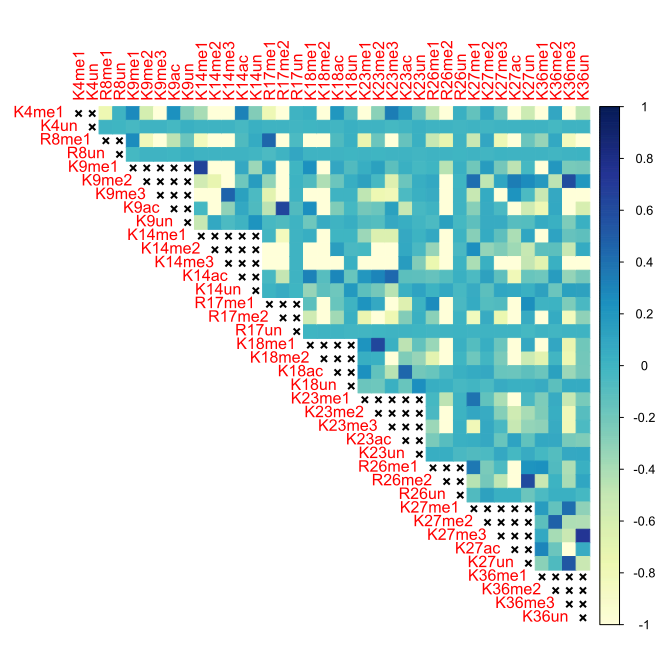

### Figure S22

Kidney, 18 months, Histone H3.2

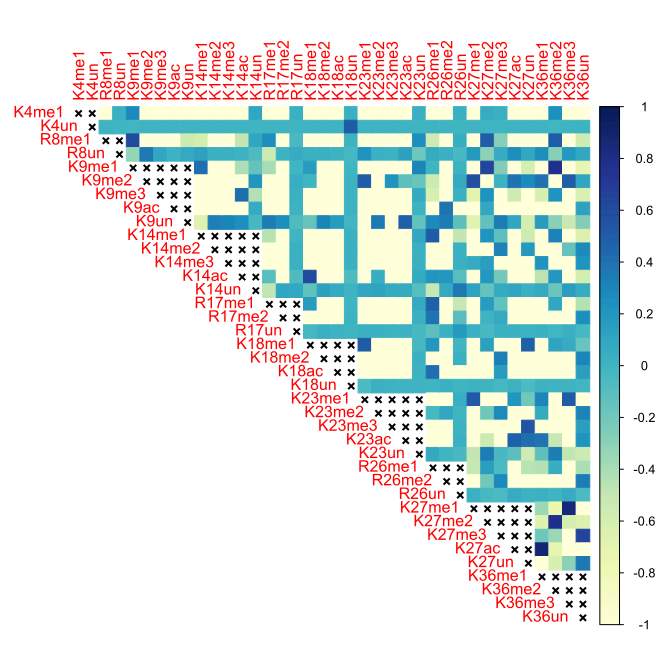

### Figure S23

Kidney, 18 months, Histone H3.3

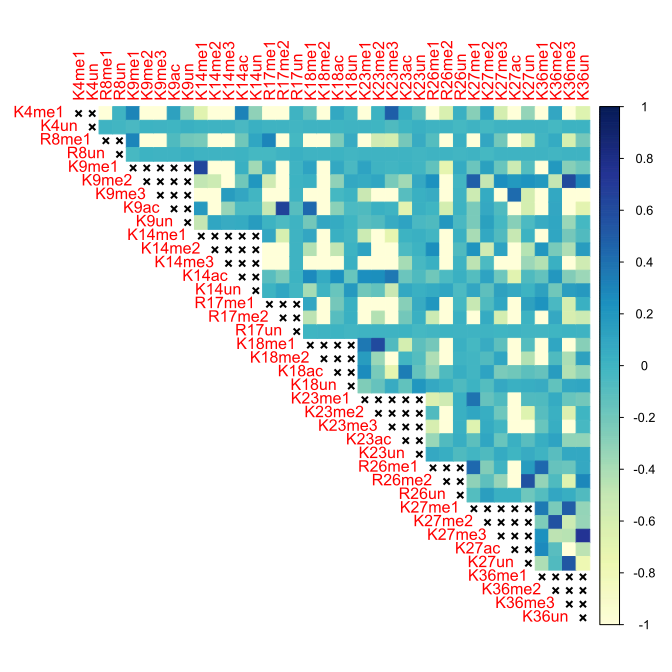

### Figure S24

Kidney, 24 months, Histone H3.2

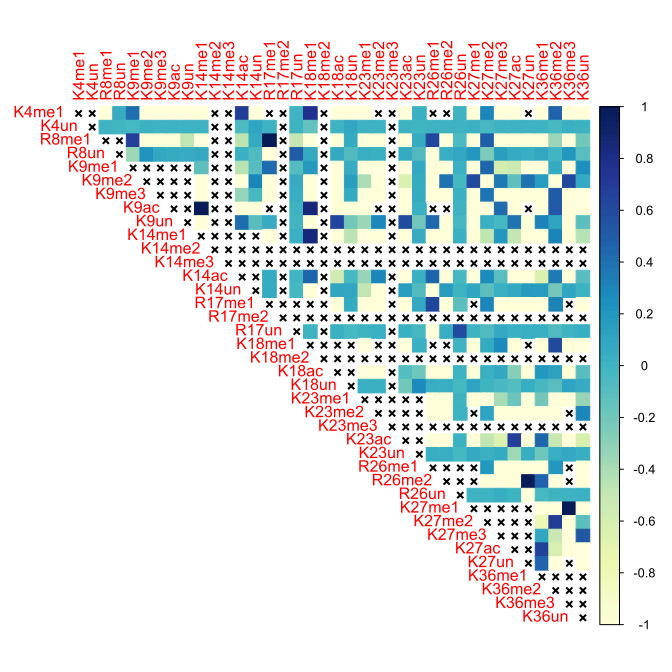

### Figure S25

Kidney, 24 months, Histone H3.3

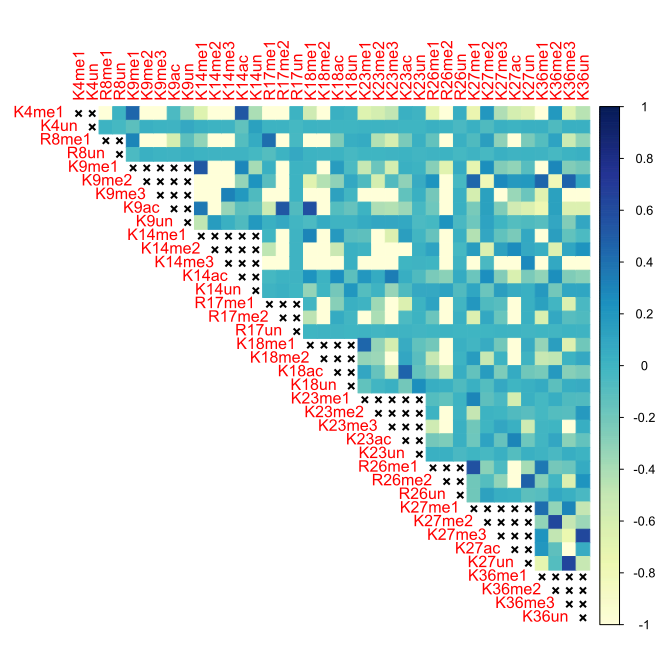

### Figure S26

Kidney, 3 months, Histone H3.2

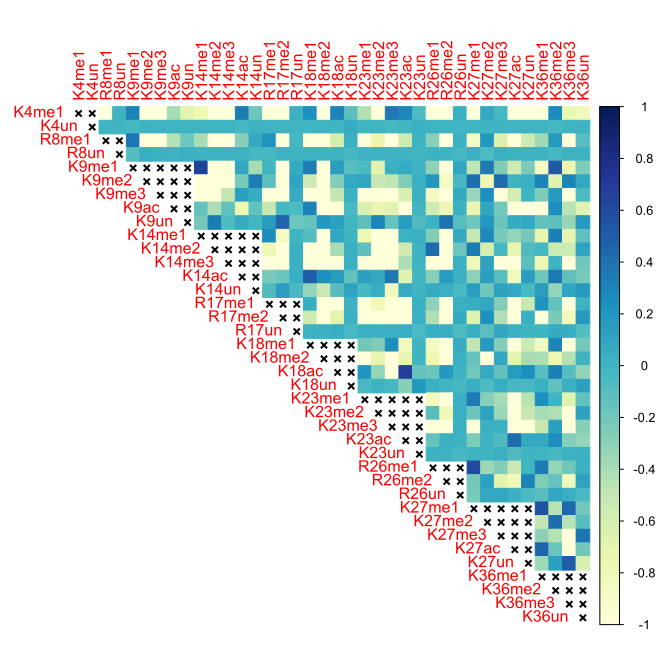

### Figure S27

Kidney, 3 months, Histone H3.3

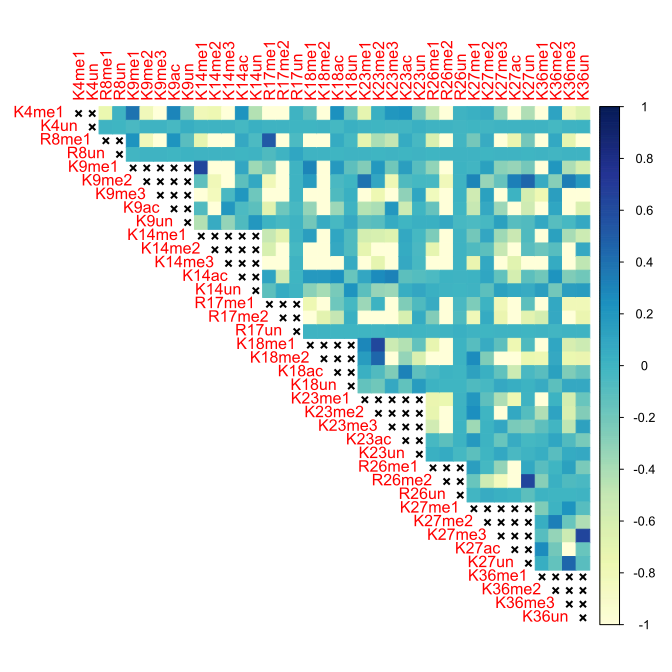

### Figure S28

Kidney, 5 months, Histone H3.2

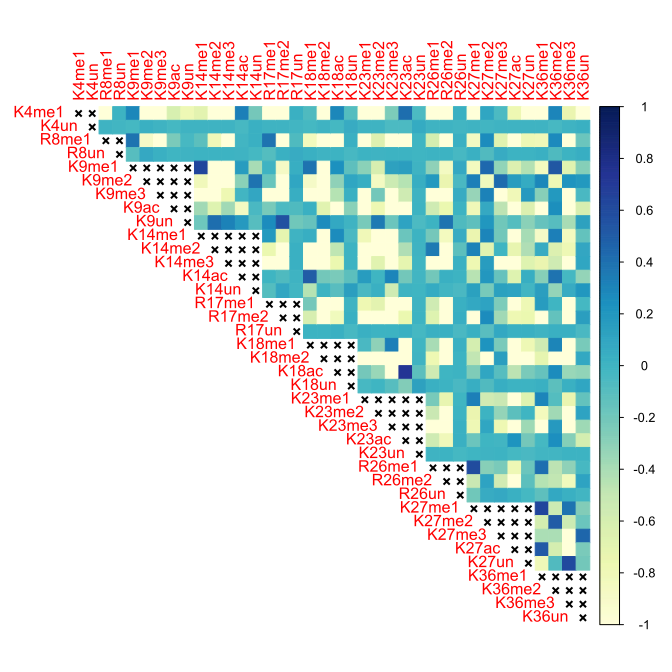

### Figure S29

Kidney, 5 months, Histone H3.3

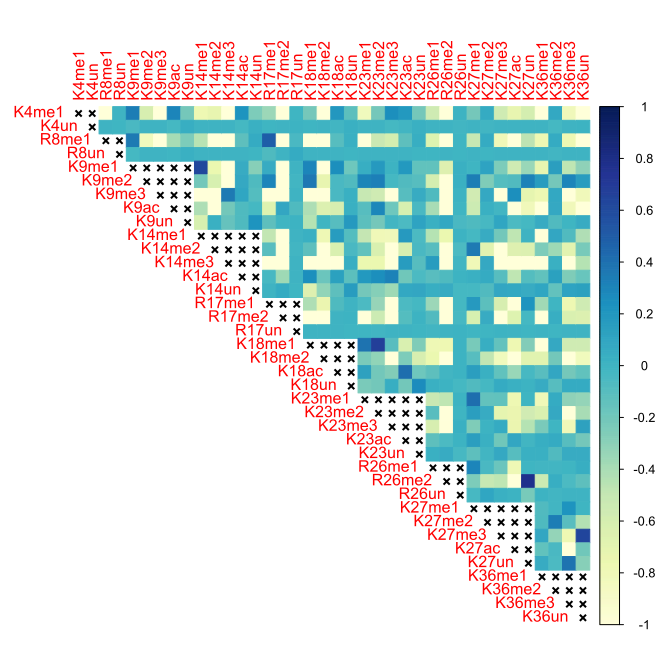

### Figure S30

Liver, 10 months, Histone H3.2

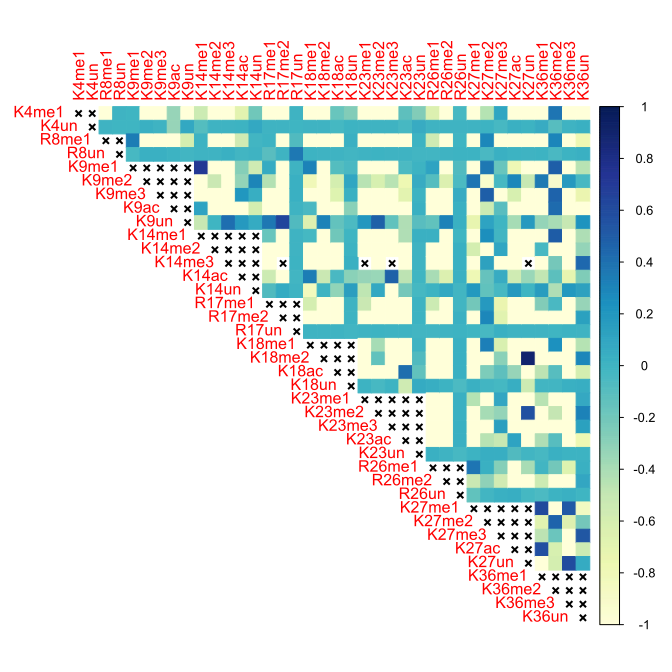

### Figure S31

Liver, 10 months, Histone H3.3

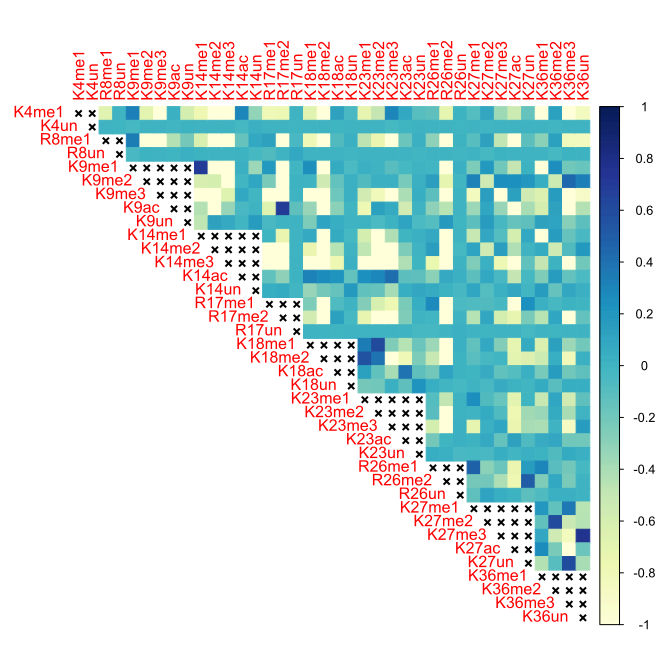

### Figure S32

Liver, 18 months, Histone H3.2

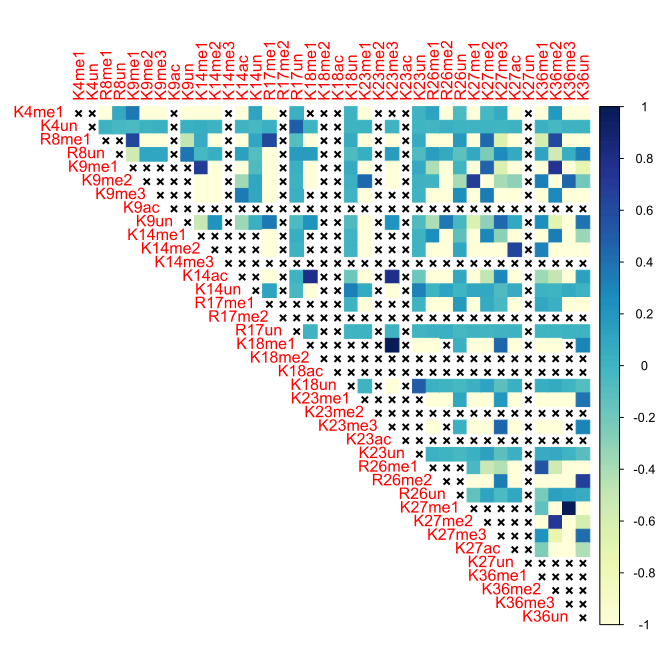

### Figure S33

Liver, 18 months, Histone H3.3

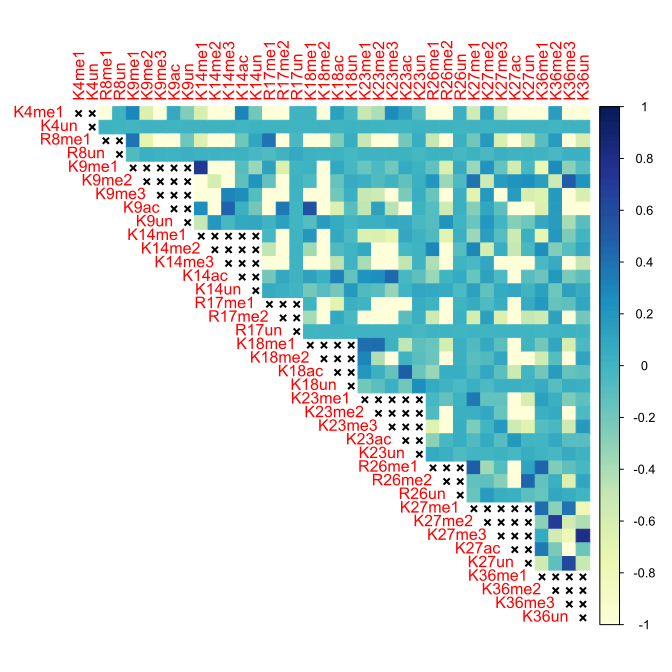

### Figure S34

Liver, 24 months, Histone H3.2

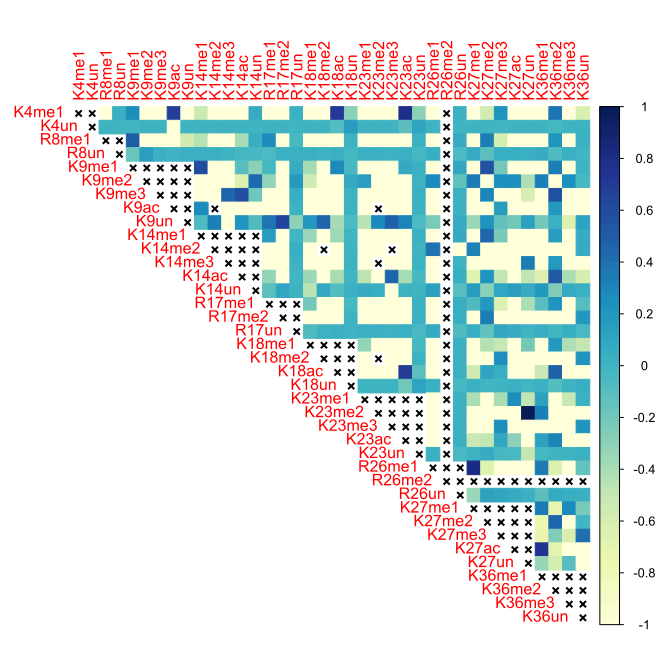

### Figure S35

Liver, 24 months, Histone H3.3

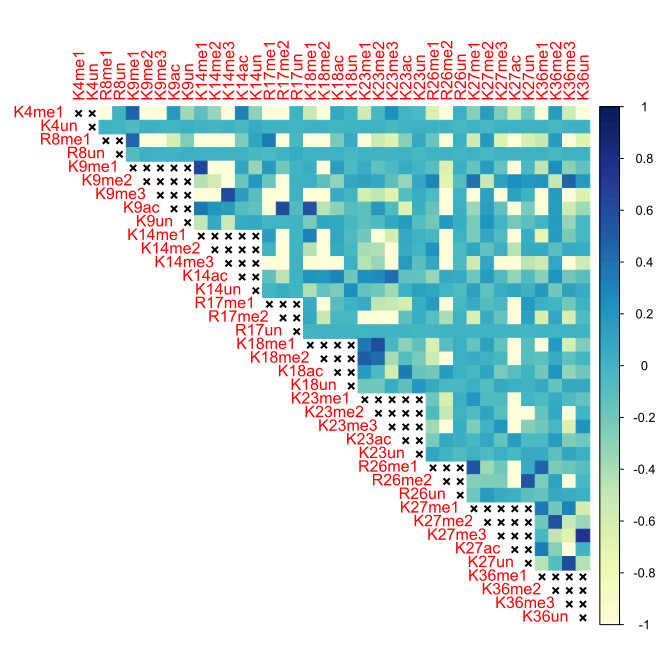

### Figure S36

Liver, 3 months, Histone H3.2

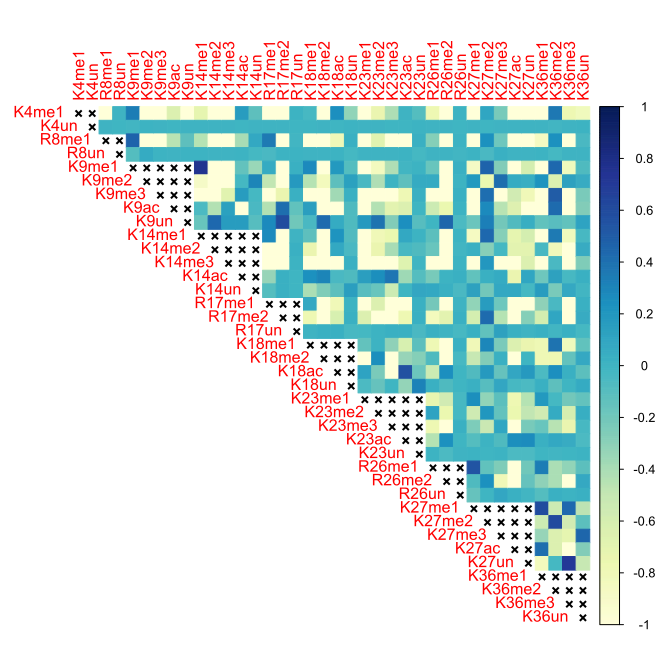

### Figure S37

Liver, 3 months, Histone H3.3

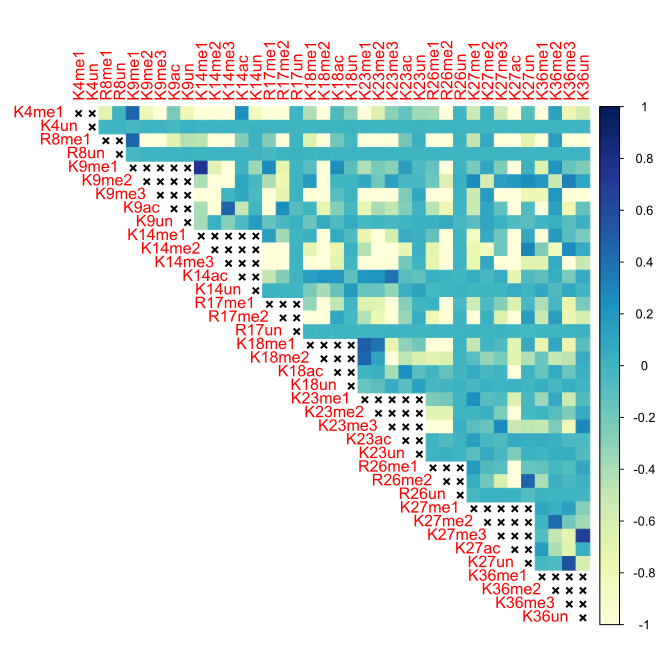

### Figure S38

Liver, 5 months, Histone H3.2

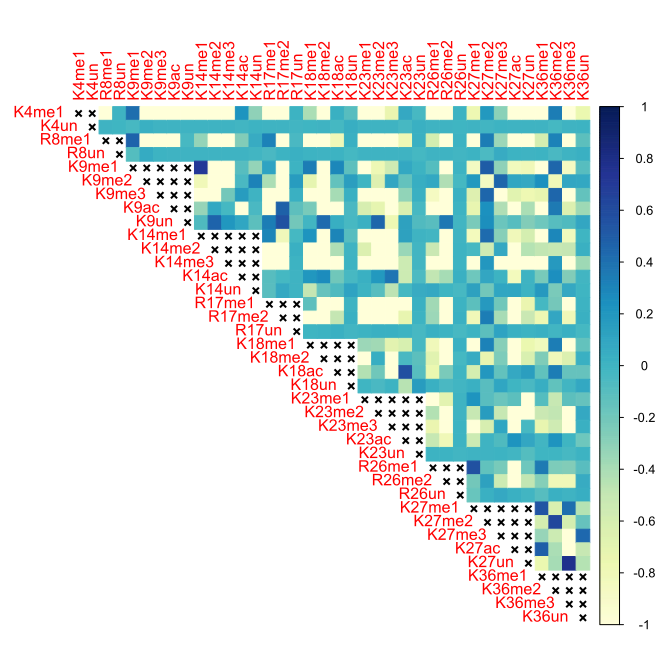

### Figure S39

Liver, 5 months, Histone H3.3

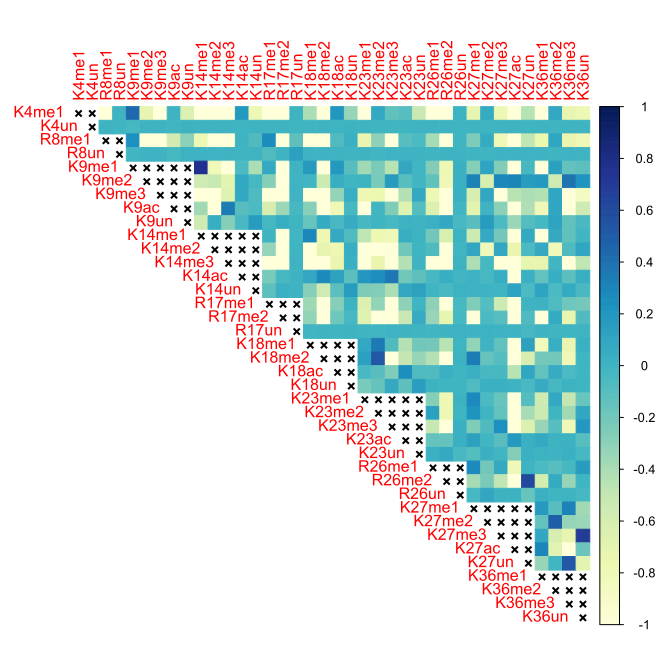
